# Environmental conditions define the energetics of bacterial dormancy and its antibiotic susceptibility

**DOI:** 10.1101/2020.06.18.160226

**Authors:** L Mancini, T Pilizota

## Abstract

Bacterial cells that stop growing but maintain viability and the capacity to regrow are termed dormant and have been shown to transiently tolerate high concentrations of antimicrobials. The proposed mechanism behind the enhanced survival capabilities of these cells is the reduced energy supply. However, not all reported results are in agreement, and the exact role of energetics remains unsolved. Because dormancy merely indicates growth arrest, which can be induced by various stimuli, we hypothesise that dormant cells may exist in a range of energetic states that depend on the environment. We first establish conditions that are capable of inducing dormancy, and subsequently measure the energy profiles they elicit in single dormant cells. Our simultaneous measurements of proton motive force (PMF), cytoplasmic pH and ATP concentrations confirm that dormant cells exhibit characteristic energetic profiles that can vary in level and dynamics, depending on the stimulus leading to growth arrest. We test whether the energetic makeup is associated with survival to antibiotics of different classes and find that, while growth arrest remains the dominant mechanism enabling survival, some correlations with cellular energetics exist. Our results pave the way to a classification of dormant states based on energy profiles, support a novel relationship between environment and drug susceptibility of dormant cells and suggest that knowledge of the conditions present at the infection site is necessary to design appropriate treatments.

## Introduction

The global surge in bacterial antibiotic resistance in both clinical and environmental isolates highlights the need for a better understanding of bacterial survival strategies. Recent work Balaban et al. [2019], Brauner et al. [2016] divides antimicrobial survival strategies to antibiotic challenges into three main classes: resistance, tolerance and persistence. Resistant bacteria carry genetic mutations that allow them to thrive at antibiotic concentrations above the Minimum Inhibitory Concentration (MIC) Brauner et al. [2016]. Tolerant bacteria may or may not carry mutations, but they only manage to withstand antibiotics for a certain amount of time Brauner et al. [2016], Levin-Reisman et al. [2017]. Persistent bacteria are genetically undistinguishable from the susceptible population and develop a phenotypical form of tolerance Balaban et al. [2019], Brauner et al. [2016].

For both tolerant and persistent cells growth cessation is often synonym of a successful survival strategy and dormancy is regarded as the silver bullet against a wide spectrum of antibiotics Lewis [2007], not only enabling infection recalcitrance Grant and Hung [2013], but also facilitating the development of resistance Levin-Reisman et al. [2017], Liu et al. [2020], Sebastian et al. [2017], Windels et al. [2019].

Microbes can enter dormancy for a number of reasons, but the mechanisms by which dormancy is attained and maintained, and how it leads to survival are multi-faceted and often poorly understood Rittershaus et al. [2013], Ronneau and Helaine [2019]. Many stimuli and environmental conditions can lead to dormancy. For example, low temperatures do so by limiting protein production Farewell and Neidhardt [1998] and folding Ferrer et al. [2003], and starvation triggers a complex signalling cascades that launch stress responses Hengge-Aronis [2002], Pletnev et al. [2015] and halt DNA replication Ferullo and Lovett [2008]. Internal stimuli, such as DNA damage Kreuzer [2013] or toxin production Coussens and Daines [2016], can also initiate growth arresting responses. Lastly, quorum sensing molecules can limit cell energy Krasnopeeva et al. [2019], Lazazzera [2000], and a range of environmental toxins can stop growth without impacting viability, and, many of these populate our arsenal of bacteriostatic antibiotics Baquero and Levin [2021].

Although dormancy can be achieved in multiple ways, these all lead to an easily observable phenotype: growth arrest. Growth and metabolism are naturally connected spurring a number of investigations into a possible correlation between cellular energy (ATP in particular) and survival capabilities of dormant cells Aedo et al. [2019], Braetz et al. [2017], Conlon et al. [2016], Leszczynska et al. [2013], Lobritz et al. [2015], Shan et al. [2017], Svenningsen et al. [2019], Wilmaerts et al. [2018]. The results are seemingly contradictory. On one hand, stationary *Staphylococcus Aureus*, arsenate-poisoned and HokB toxin expressing *Escherichia coli* were found to have reduced levels of ATP and high tolerance to ciprofloxacin and ampicillin and ofloxacin, respectively Conlon et al. [2016], Shan et al. [2017], Wilmaerts et al. [2018]. Similarly, *E. coli* cells with high ATP levels were found more susceptible to *β*-lactams and fluoroquinolones Aedo et al. [2019] and cells with low respiration rates were found to deal better with several bactericidal antibiotics Lobritz et al. [2015]. On the other hand, *Salmonella enterica* cells with reduced ATP levels due to deletion of the *atp* operon were more susceptible to ciprofloxacin Braetz et al. [2017] and ATP concentration was not low in ampicillin-tolerant stationary phase and amino-acid starved *E. coli* Leszczynska et al. [2013], Svenningsen et al. [2019]. These apparently contradictory results could be reconciled if dormant states had different energy levels (including different temporal profiles of energy levels), and hence drug susceptibilities. As this would have relevant repercussions on treatment design, we systematically mapped, at the single-cell level, the energetics of growth arrested cells that were dormant in response to different external stimuli. Specifically, we measured the cellular electrochemical gradient of protons (proton motive force, PMF), its ΔpH component across the plasma membrane and the ATP concentration. PMF plays a fundamental role in cell physiology, powering transport across the plasma membrane Bradbeer [1993], Jahreis et al. [2008], Ramos and Kaback [1977], Wood [2015], including those mediated by multi-drug-efflux pumps Anes et al. [2015], and fuelling the production of ATP via F_1_F_*o*_ ATP synthase. The latter can reversibly transform ATP and PMF into one another Mitchell [1961]. The ΔpH component of the PMF is directly linked to the intracellular pH, which is maintained near neutral in organisms across all kingdoms of life. Recently, it was discovered that the ability to maintain intracellular pH at different extracellular pHs depends on the absolute value of the PMF itself Terradot et al. [2021].

Our results show that the energetic states of cells that entered dormancy due to different extracellular signals are not equivalent, and can range from growth-like to markedly reduced. In addition, we found that the energy profile and levels, characteristic for a given dormancy-inducing stimulus, can influence the survival to antibiotics of different classes in an energy-dependent manner. The importance of energetics could, to a first order approximation, be explained with the role of PMF/ATP on characteristic antibiotic accumulation. Our work portrays dormancy as a phenotypically complex state, a notion that we expect applies to similar states in other organisms, and that shows that knowledge of the environmental conditions present at the infection site is important for the design of accurate treatment strategies.

## RESULTS

### Carbon starvation and bacteriostatic drugs can induce dormancy instantaneously

To investigate the energetics of bacterial dormancy we focused on *E. coli* and started by establishing a working definition of dormancy based on its most widely accepted hallmarks: growth arrest and viability. We note that this is one requirement less than Gefen et al. [2008], where in addition to being viable and non-growing, dormant cells also did not produce a reporter protein (interpreted as absence of protein production as a whole). To start, we tested six different conditions that bacteria can encounter during infection and that could cause dormancy: (a) carbon starvation, (b) nitrogen starvation, (c) indole (a putative quorum sensing molecule produced by stationary phase cultures that has been shown to act as a protonophore Chimerel et al. [2012], Kim and Park [2015], Krasnopeeva et al. [2019]), (d) chloramphenicol treatment (bacteriostatic antibiotic that stops protein synthesis Gale and Folkes [1953]), (e) rifampicin treatment (bacteriostatic antibiotic blocking RNA synthesis Hartmann et al. [1967]) and (f) trimethoprim treatment (bacteriostatic antibiotic that halts DNA synthesis Brogden et al. [1982]). We assayed whether these arrest growth immediately upon addition, and evaluated post-treatment viability. To assess the growth, we grew the cells in exponential phase for ~ 15 generations and then either moved them to the same growth media but with no carbon or nitrogen, or exposed them to different concentrations of the bacteriostatic drugs or indole (Fig. 1A and SI Fig. 1). To estimate growth we monitored bacterial cell numbers over time via optical density (OD). Fig. 1A shows that carbon starvation, 200 *μ*g/ml chloramphenicol, 16*μ*g/ml rifampicin and 0.2*μ*g/ml trimethoprim cause immediate growth arrest, as indicated by the fact that OD remained constant. However, OD is an indirect measure of cell number, with a proportionality that is linear only as long as cell size and index of refraction remain constant Stevenson et al. [2016]. Thus, we checked that our candidate dormancies were not causing significant cell size changes by monitoring the cells in a microscope. Via microscopy we also confirmed that constant OD was not a result of matching growth and lysis rate. Removing just the nitrogen source did not cause growth arrest, perhaps, and as pinpointed by others before Kim et al. [2012], because traces of ammonium are often found in various chemicals and can, at least to some extents, support bacterial growth (SI Fig. 1A). Treatment with indole stopped cell growth at a concentration of 10 mM (SI Fig. 1B). We next tested cells’ viability under our candidate dormancies by withdrawing aliquots from the cultures after 30, 60, 120, 180 and 240 min of treatment, washing them into fresh media and counting the colony forming units (CFU), Fig. 1B. We found that the chloramphenicol decreased viability to half over the 4 h while starvation rifampicin and trimethoprim treatment had no effect on cell viability. Lastly, indole did not satisfy our definition of dormancy because at 10 mM it was lethal to 99.7% of the cells (SI Fig. 1F), and smaller concentrations of indole did not arrest growth.

**Figure 1.**
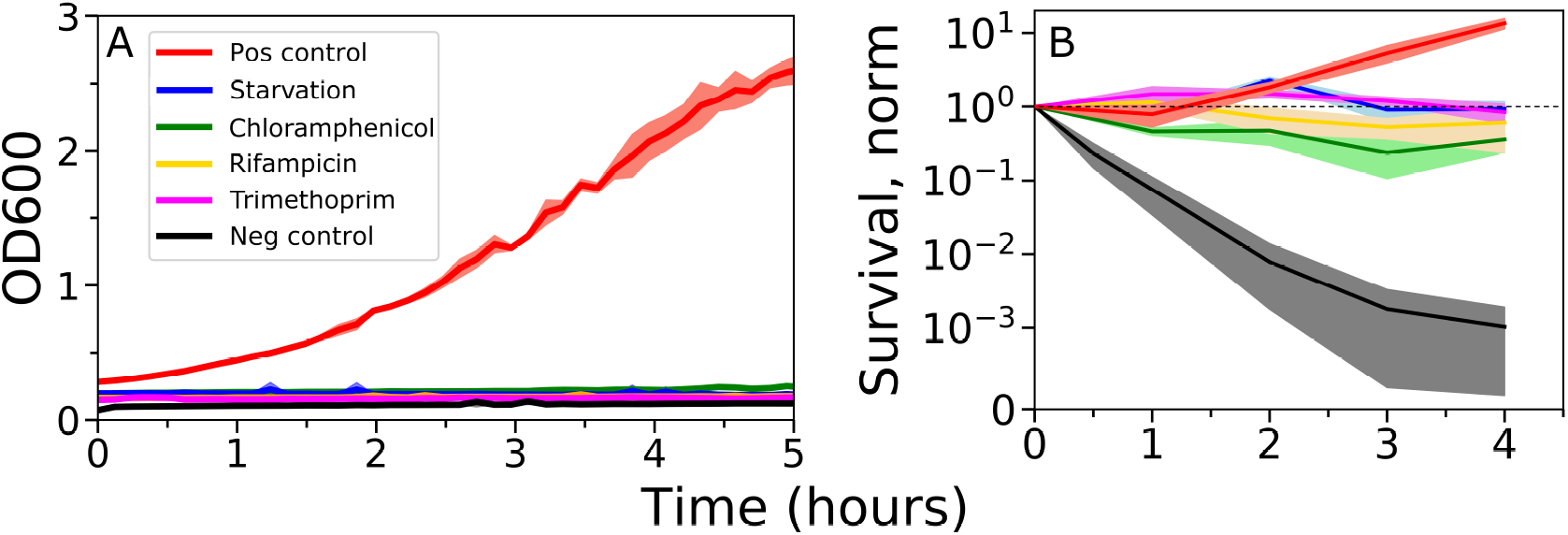
Individuation of different dormancies by monitoring cell growth and viability. **a,** addition of 200 *μ*g/ml of chloramphenicol, 16 *μ*g/ml rifampicin, 0.2 *μ*g/ml trimethoprim or removal of the carbon source sharply arrest growth of previously exponentially growing cells. The shaded area shows the standard deviation of 3 replicates and experiments were performed in a plate reader. **b,** Cell viability under candidate dormancies identified in **a** assayed by counting colony forming units (CFUs). The error bars show the standard deviation of 3 independent replicates. Treatment with 5xMIC ciprofloxacin, a bactericidal antibiotic, is given as a negative control, and untreated cells as a positive control. The dashed line indicates constant cell number.

### The PMF of dormant cells varies in a condition-dependent manner

Having individuated dormancies that satisfied our definition, we investigated their energy levels by measuring single cell PMF and one of its two components, ΔpH across the plasma membrane, using assays we previously developed Krasnopeeva et al. [2019], Mancini et al. [2020], Wang et al. [2019]. Briefly, we used bacterial flagellar motor speed as a proxy for relative PMF changes because it varies linearly with PMF Fung and Berg [1995], Gabel and Berg [2003]. To measure the motor speed we attached bacterial cells to the surface of a glass coverslip and polystyrene beads to genetically modified, truncated flagellar filaments Krasnopeeva et al. [2019], Rosko et al. [2017], Ryu et al. [2000], Wang et al. [2019]. We measured the bead position, now rotated by the motor, with back focal plane interferometry thus gaining information on motor speed Krasnopeeva et al. [2019], Rosko et al. [2017], and we we carried out the measurements continuously throughout the treatment to include the transition into dormancy as well as the energy levels during dormancy. The method allows us to measure the PMF of one cell per experiment (the one placed in the heavily attenuated optical trap). To simultaneously measure ΔpH we expressed ratiometric, pH sensitive fluorescent protein in the cell cytoplasm and read its ratiometric signal every 90 s. We did so for the cell in which we are simultaneously measuring the PMF, but also on all other neighbouring cells in the same field of view, Fig. 2A. Illumination conditions used have been previously characterised – these resulted in negligible photobleaching and caused no damage to *E. coli* Krasnopeeva et al. [2019]. SI Fig. 2 shows the calibration curve we used to convert ratiometric measurements to pH Wang et al. [2019]. Finally, we also measured the extracellular pH. We performed our measurements on cultures that had grown for ~ 15 generations in fresh medium (SI Fig. 3), therefore reaching balanced exponential growth with a doubling time of 57 min, before transition into dormancy Harvey and Koch [1980], Maaløe [1966]. Fig. 2 shows characteristic energy profiles of different dormancies. In particular, cells treated with chloramphenicol lost only *~* 25% of their PMF over the first 20 min of treatment settling onto this baseline thereafter (Fig. 2D, green trace). Cells treated with rifampicin did not lose PMF over the first 30 min, but after ~ 2 h the PMF was ~ 30% lower. Cells treated with trimethoprim gradually lost ~ 15% of their PMF over 30 min and we could not find any spinners after 2 h treatment. We could not establish whether this was due to full loss of PMF or whether the drug could have any effects on the expression of motors. Cells experiencing carbon starvation settled onto a new, ~ 50% lower PMF baseline within 2-3 min (Fig. 2D, blue trace). In all cases, internal pH (Fig. 2B) decreased as well. By the end of the period we measured (10 or 30 minutes), it dropped by ΔpH 0.35, 0.2, 0.25 and 0.8 for chloramphenicol, rifampicin, trimethoprim and carbon starvation, respectively.

**Figure 2.**
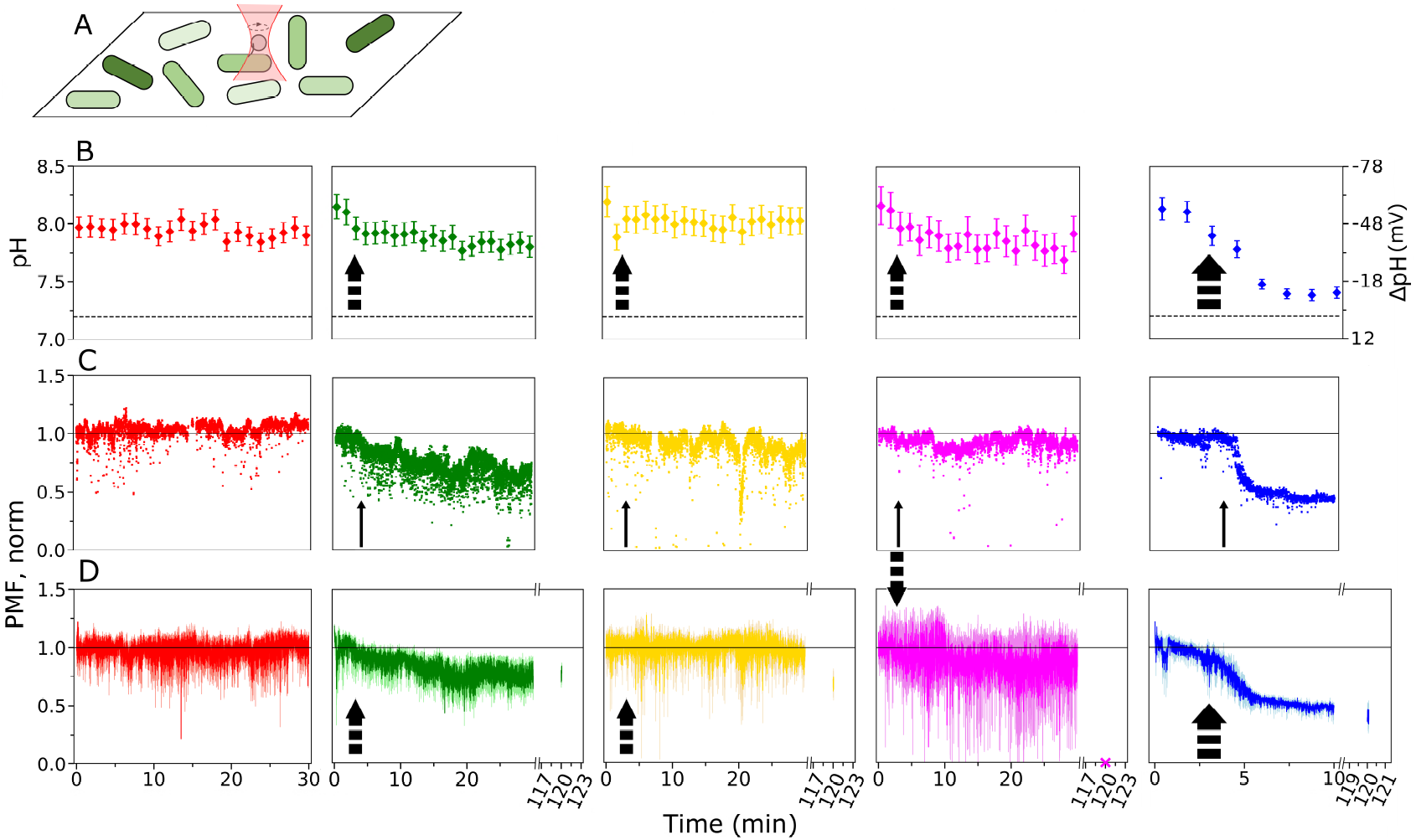
Different dormancies are not energetically equivalent. **a,** Cartoon of a typical field of view in which the flagellar motor speed of a cell is measured simultaneously as its cytoplasmic pH. The pH of all of the neighbouring cells in the field of view is also measured. **b,** Average cytoplasmic pH and ΔpH for cells in growth medium (M63 with glucose shown in red as a negative control), with 200 *μ*g chloramphenicol (green), 16 *μ*g rifampicin (yellow), 0.2 *μ*g trimethoprim (fuchsia) or medium deprived of the carbon source (blue). Fresh media is continuously supplied to the cells and error bars are standard errors. Average values for each time point were obtained from 8 to 10 experiments for a total of 494 cells in M63 with glucose (media), 605 cells in the media with chloramphenicol, 796 cells in the media with rifampicin, 593 cells in the media with trimethoprim and 594 cells in media with no glucose. Due to our flow cell design [Mancini et al., 2020] the switch between growth medium used during slide preparation and dormancy-inducing medium occurs 2 to 4 min (black arrow, where the width of the arrow indicates the duration of the interval) from the beginning of the recording. The dashed line indicates the pH of the medium, which is 7.2. **c,** Example single-cell, normalized PMF measurements from individual motor speed traces, each in one of the five conditions shown in **b**. **d,** Average normalized PMF of 10 cells in each condition. Standard errors are given. In the case of carbon starvation, chloramphenicol and rifampicin treatment, we also assayed the PMF after 2 h incubation on 3, 24 cells and 6 cells, respectively. It was not possible to find spinning motors after 2 h in the trimethoprim condition, which could indicate that the PMF may have been lost, we indicate this with a cross. To adjust for the variable arrival time of the treatment to the field of view (after 2-4 minutes), the 10 PMF traces of the carbon starvation condition (blue) were aligned as explained in *Methods*.

### An improved QUEEN-based ATP sensor reveals ATP and PMF coupling during dormancy

Having measured PMF levels during dormancy we next focused on the ATP concentrations. Because we wished to measure time courses, ideally concurrently with single-cell PMF measurements, we chose the QUEEN ATP sensor Yaginuma et al. [2014]. However, the sensor suffers from poor signal intensity and requires exposure times of 1-2 s at short wavelengths, which can quickly lead to cell damage Krasnopeeva et al. [2019], Mancini et al. [2020]. Therefore, we focused first on optimising the sensor’s expression level and signal-to-noise ratio. To improve the expression level we placed the sensor (QUEEN 2mM and QUEEN 7*μ*M Yaginuma et al. [2014]) downstream of the strong constitutive promoter of the cytochrome C oxidase from *V. Harveij* Pilizota and Shaevitz [2012]. We also removed the histidine sequence with which the QUEEN sensors have been tagged in Yaginuma et al. [2014] for protein purification finding that, particularly for QUEEN 7*μ*M, the removal of the tag produced an increase in signal intensity (Fig. 3A, SI Fig. 4A). We named these brighter derivatives QUEEN 2mM* (Q2*) and QUEEN 7*μ*M* (Q7*). Because in Yaginuma et al. [2014] the His-tag is positioned in the proximity of the ATP sensing region we reasoned that its removal could have an effect on the protein’s response to ATP (especially because *in vivo* ATP is commonly found in coordination with magnesium ions Gout et al. [2014]). Thus, we characterized Q7*, the brightest sensor, *in vitro*, working with cells’ lysates (see *Methods*). The removal of the His-tag did not change spectral properties of the protein (SI Fig. 4B), but the sensitivity range shifted towards higher [ATP] (Fig. 3B), making Q7* suitable for performing ATP measurements in the physiological range of *E. coli* Yaginuma et al. [2014]. We also found that the sensor exhibits a slight temperature dependence, shifting the sensitivity range toward higher [ATP] at 37°C compared to room temperature (SI Fig. 4E). The calibration should, therefore, be performed for the specific temperatures at which the experiments are carried out. We next tested the pH sensitivity of Q7*. We found that the sensitivity of the sensor to [ATP] changes little in the pH 7.2 to 7.6 range. For pH above 7.8 the fluorescent ratio (of the 400/488 nm) decreases for the same [ATP], and is overall significantly less sensitive to [ATP] changes. Next, we assessed the sensor’s photostability, first focusing on whether photobleaching altered ATP calibration (SI Fig. 5). Again, using cell lysates for the *in vitro* characterization, we observed that both the 400 and 488 nm wavelengths can photobleach (SI Fig. 5A-C). We also characterized the sensor’s photostability inside the cells. Because short wavelength illumination can lead to significant photodamage Krasnopeeva et al. [2019], we monitored bacterial PMF under different illumination frame rates. We found that when illuminated every 7 min with 488 and 400 nm wavelengths for 20 min total, cells experience no photodamage (Fig. SI 5D-F). We then sought to decouple the behaviour of the QUEEN sensor from cellular physiology by treating bacteria with 5% ethanol, which drops the PMF Krasnopeeva et al. [2019] and presumably limits any physiologically driven changes in the intracellular [ATP]. After 15 min, and still keeping the ehanol in, we exposed the cells to illumination at different frame rates. As expected, we observed exposure-dependent photobleaching at the 400 nm wavelength (SI Fig. 5G). Contrary to our observations with lysates, the signal from the 488 nm wavelength showed a time-dependent intensity increase (SI Fig. 5H), where the 400/488 ratio decreased significantly with illumination (SI Fig. 5I). The time-dependent increase in intensity under 488 nm illumination could be explained by photoactivation, a light-dependent red shift common among GFP and its derivatives Elowitz et al. [1997], Patterson and Lippincott-Schwartz [2002], or by a drop in the [ATP]. As the cells have been pre-treated with ethanol, we deemed the latter less likely, but we further confirmed it by exposing cells to even stronger concentrations of PMF decouplers: a mixture of 2% ethanol and 10 mM indole which acts as a protonophore and immediately shuts down PMF (SI Fig. 6C) Chimerel et al. [2012], Kim and Park [2015], Krasnopeeva et al. [2019]. In SI Fig. 6 we show that in these cells we observe similar behaviour to that in SI Fig. 5. We concluded that the increase in intensity observed under 488 nm illumination was most likely caused by photoactivation, and that given the sensor’s photoproperties, time series measurements should be replaced with an estimate of cellular [ATP] at only one time point. In dormant cells we choose for that point 10 min after initiation of dormancy for carbon-starved cells and 30 min for chloramphenicol treated cells. We did not measure [ATP] in the other two bacteriostatic drugs, because these only caused small PMF changes in the 30 min interval. As a negative control we measured [ATP] in the cells prior to treatment. And, as a positive control, we measured [ATP] in cells treated with the 2% ethanol+10 mM indole mixture (after 10 min of treatment). Because our pH and ATP sensors share similar spectral properties, pH and ATP measurements could not be performed in the same cells. Instead, we carried out [ATP] measurements while simultaneously monitoring flagellar motor speed, in a similar fashion to Fig. 2 (where we simultaneously monitored PMF and pH). The trends of PMF upon treatment for strains expressing the Q7* ATP sensor recapitulated the ones we observed in Fig. 2. As shown in Fig. 3C, cells treated with a bacteriostatic antibiotic maintain seemingly constant ATP levels, whereas cells deprived of carbon quickly lose one third of their ATP, in good correlation with the PMF trends. As expected, cells treated with the ethanol and indole mixture show the most marked ATP loss. The [ATP] measurements were done at the following cytoplasmic pHs: 8.1 for the positive control, 7.9 for chloramphenicol, 7.4 for carbon starvation and 7.2 for the negative control. At pHs above 7.8 the 400/488 nm ratio of the Q7* sensor decreases (SI Fig. 4E), thus the ATP values for untreated and antibiotic-treated cells may be underestimated. However, they still confirm the overall trend of cellular energetics. The observed ATP concentrations are in the same order of magnitude of those reported by Yaginuma et al. [2014].

**Figure 3.**
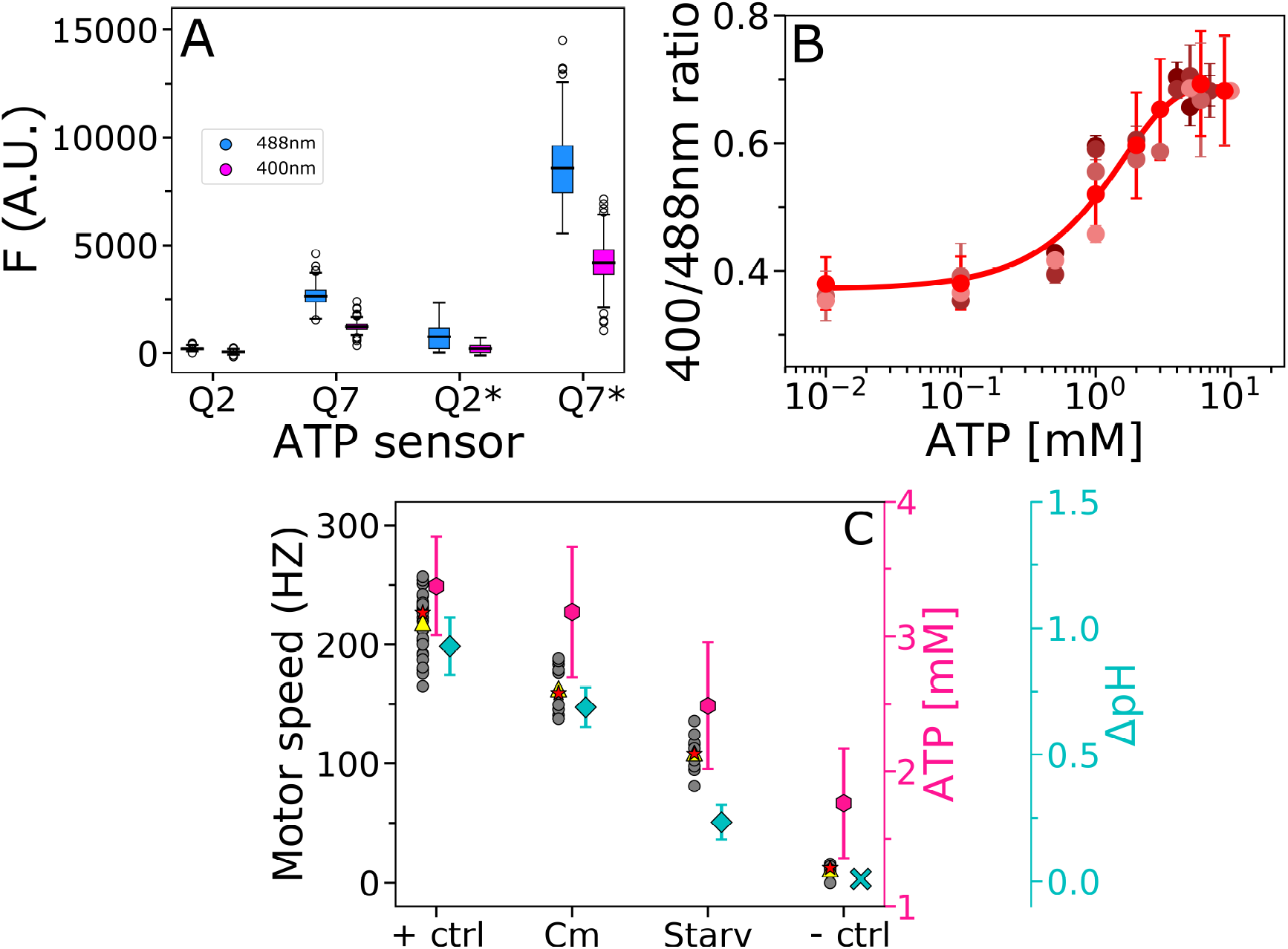
QUEEN 7*μ*M* shows improved signal intensity and can sense in *E. coli*’s physiological range. **a,** Comparison of the signal intensity from the four sensors. Fluorescence values for the two wavelengths of interest for the QUEEN sensor (400 and 488 nm) were obtained from 270 single cells expressing pWR20-Q2, 259 expressing pWR20-Q7, 194 expressing pWR20-Q2* and 315 expressing pWR20-Q7*. All cells were imaged in a tunnel slide (designed as given in Rosko et al. [2017]), with the same exposure time and gain (see *Methods*). The box plot shows lower and upper quartile and median value, with the whiskers indicating lowest and highest extremes.**b,** Calibration curve of the Q7* sensor at 37°C and pH 7.7. Each data point is the average of 3 or more replicates, the error bars show the standard deviation. Different shades of red indicate 5 independent experiments (each performed in replicates). Data from experiments performed on different days were re-scaled as explained in the *Methods* section. The fit (red line) to all of the data points is used for determining ATP concentration in (**c**). **c,** ATP and ΔpH of cells in different dormancies measured with QUEEN 7*μ*M* sensor. Measurements were made after 30 (negative control and chloramphenicol) and 10 min (carbon starvation and positive control) from the beginning of the treatment. Grey circles indicate individual motor speeds under different conditions (median of 5 s long recording for each cell). Total of 20, 11, 10 and 10 cells for negative, chloramphenicol-treated, carbon starved and positive control cells, respectively, are given. Red stars indicate the mean, and yellow triangles the median values. The cytoplasmic pH values used to calculate ΔpH (cyan) are from the experiments in Fig. 2A. Because indole interacts with our pH sensor Wang et al. [2019] we could not measure it, but assumed ΔpH of cells treated with indole is zero because indole is a protonophore Krasnopeeva et al. [2019]. We indicate this value with a cyan cross. ATP concentration (pink) was measured in 493 chloramphenicol treated and 633 carbon starved cells. For positive control we measured [ATP] of 1176, and for negative control 793 individual cells. The error bars show the standard error.

### Dormant states with different energy levels show various degrees of susceptibility to antibiotics of different classes

Having established that different dormancies have different energetic profiles, we wondered if these influenced antibiotic susceptibility. Previous findings show that cellular energy levels play a role in antibiotic efficacy, e.g. by governing aminoglycoside drugs uptake rate Taber et al. [1987], fuelling multi-drug efflux pumps activity Anes et al. [2015] or other cellular response mechanisms such as DNA repair Kowalczykowski [1991], Kowalczykowski et al. [1994]. We exposed exponentially growing cells to the four dormancies, as in Fig. 2. After 30 min, when cells were dormant, we exposed the cultures to 5x MIC concentration of two bactericidal drugs: ciprofloxacin, a DNA damaging drug thought to be a substrate of multi-drug efflux pumps Piddock [2006] and kanamycin, an aminoglicoside that causes errors in protein translation Jelenc and Kurland [1984]. We point out that cells were kept in dormancy-inducing conditions during the treatment with bactericidal drugs. The MIC concentrations were obtained as explained in *Methods* and shown in SI Fig. 7. We assayed survival of dormant cells under ciprofloxacin and kanamycin treatment for up to 4 h, by withdrawing aliquots at regular intervals and performing CFU counting. Fig. 4A illustrates our experimental protocol. For all dormancies tested, cells cope significantly better with antibiotic challenges than fast growing cells (positive control), confirming the protective role of dormancy, Fig. 4B and C. Overall, the dormancies we selected offered better protection from kanamycin treatment (Fig. 4B-C), but we also observed dormancy-specific differences. For both ciprofloxacin and kanamycin, we noticed a correlation between PMF and the level of dormancy protection, Fig. 4D and E. In the presence of ciprofloxacin (Fig. 4D), carbon starved cells died faster than cells treated with bacteriostatic antibiotics, with only 3 cells in 1000 remaining alive after 3 h of treatment. Chloramphenicol-treated cells died the slowest; even after 4 h more than a fourth of the population was still alive. In the case of kanamycin (Fig. 4E), dormancy-specific differences were small, but carbon-starved cells, which have the lowest PMF, seemed to fare better against the drug. Ciprofloxacin is a known substrate of PMF-dependent efflux pumps in *E. coli*, while aminoglycosides, like kanamycin, are known to accumulate in highly energetic cells. Our results could therefore be coherent with an energy-dependent model of accumulation of the bacteriolytic antibiotics whereby highly energetic dormant cells deal better with ciprofloxacin, because they can proficiently use energy-fuelled efflux pumps to get rid of the drug, Fig 4D. Vice versa, scarcely energetic dormant cells, would take up less kanamycin and survive better, Fig. 4E.

**Figure 4.**
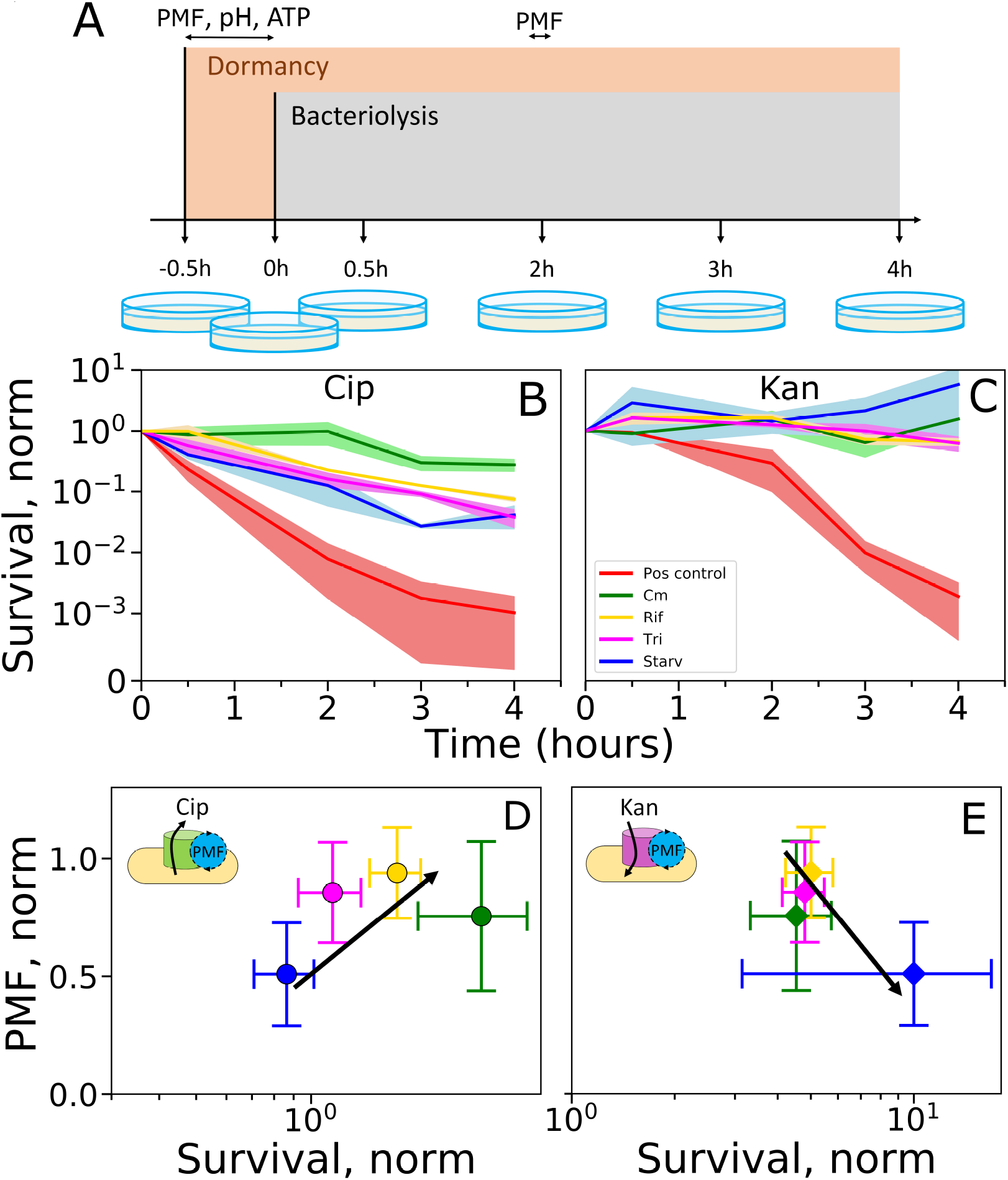
Dormant cells survival to antibiotic treatment (at 5x MIC). **a,** Cartoon showing the time points at which dormancy is induced and bactericidal antibiotics are added (orange and grey), as well as time points at which survival is estimated (petri dishes) and at which energetics are measured. The bacteriolytic antibiotics were added 30 min from initiation of dormancy, and CFUs were counted 30, 120, 180 and 240 min after the addition of the bacteriolytic antibiotics. **b,** Survival to treatment with ciprofloxacin and **c,** kanamycin. Cells that have gone dormant as a consequence of chloramphenicol, rifampicin, trimethoprim treatment or carbon starvation are shown in green, yellow, fuchsia and blue, respectively. These are compared to a positive control in which cells are not dormant. CFU counts are normalized by the value right before the bacteriolytic antibiotic addition, i.e. 30 min after initiation of dormancy. Solid lines show the median of 3 independent replicates and shaded areas the standard error of the mean. **d** and **e,** survival to antibiotic treatment plotted against normalized PMF levels of dormant cells prior to the treatment. Error bars show the standard deviation. Survival is estimated as the total area under the curves in **b,** and **c,** and normalized to to the CFU value right before addition of bacteriolytic antibiotic. The PMF is taken as the average of the last minute of the bacterial flagellar motor traces in Fig. 2C (after 29, 29, 29 and 9 min from the beginning of treatment for cloramphenicol, rifampicin, trimethoprim and starvation, respectively) and normalized to the average of the first minute.

## DISCUSSION

Many antibiotics interfere with molecular targets that are essential during cell growth. Dormancy is thought to protect bacteria by making these drug targets non-essential. Yet the picture is probably more complex as not all non-growing cells are equally protected from antibiotics Gefen et al. [2008], and the mechanisms behind these differences remain an open question. In this work, we deployed an ATP concentration sensor, a cytoplasmic pH sensor and an assay for PMF measurements, to investigate the physiology of dormant cells at the single cell level. Rather than using an approach in which dormancy, and thus drug survival, emerge stochastically at a low frequency, we chose to induce dormancy *a priori*. This had several advantages. First, it allowed us to bypass the use of high-persistent genetic backgrounds like *hipA7*, and allowing us to work in an effectively wild type background. Second, it gave us the freedom of testing various controlled conditions that can potentially occur at the infection site and directly link them to their effects on cells’ phenotype. Third, because the whole cell population is exposed to dormancy-inducing conditions, we could carry out low yield assays such as flagellar motor speed measurements yet obtain sufficient sample size. Our results show that cells that have gone dormant due to different external conditions settle onto markedly different energy profiles. In particular, across the conditions tested, energy levels range from growth-like to significantly reduced, demonstrating the existence of energy deprived dormant cells, but that not all dormant cells are low in energy. Our results also indicate that during dormancy, ATP and PMF levels are coupled.

Looking at susceptibility to different antibiotic we found several interesting points. Firstly, we could confirm that dormant cells outperform growing ones in terms of survival. Secondly, we presented evidence that for the aminoglycoside kanamycin growth arrest produces an almost complete protection, survival to the fluoroquinolone ciprofloxacin correlates with the PMF levels that are a consequence of different, dormancy-inducing environmental conditions. Intriguingly, cells treated with bacteriostatic drugs, which displayed only a mild reduction in the PMF, are almost unaffected by the antibiotic. Cells starved for carbon, that have suffered significant (50%) energy loss, perform an order of magnitude worse than their more energetic counterparts. Based on this correlation, we can speculate that energy profile during dormancy could be an important factor in survival to ciprofloxacin. This is not only in agreement with previous work showing that stationary phase cells are susceptible to the drug Zeiler [1985], but it is also coherent with reports indicating that the main mechanisms of survival to ciprofloxacin involve drug-efflux pumps and the SOS response Tran et al. [2016], both energy-costly processes Blanco et al. [2016], Händel et al. [2013]. In the case of kanamycin the protection caused by growth arrest seems to mask most energy-dependent effects in our conditions. However, a subtle trend that favours low energy cells seems to emerge. This is coherent with theories of membrane voltage-dependent aminoglycosides uptake Taber et al. [1987]. It is not excluded that more prominent trends might emerge at higher drug concentrations or in dormant conditions that are less energetic than the carbon starvation we tested.

Our results pave the way for a typologisation of dormant states depending on the characteristic energy profiles. We show that the different dormancy types are physiologically and clinically relevant, as we demonstrate correlations between environmental conditions and drug susceptibility of dormant cells. These results offer a mechanistic explanation for the different and seemingly contradictory antibiotic efficacies observed by others who have attempted to establish a general link between dormancy, energetics and treatment outcomes. Furthermore, our data showing that in some cases high energy can reduce antibiotic efficacy suggest that treatments based on the awakening of dormant cells by boosting metabolism may in some cases be counterproductive. While our evidence was obtained on cell populations that have been made dormant en masse, we expect our conclusions to be relevant for variations in energy levels that emerge stochastically across cell populations. As such we speculate that our results could be informative for dormancy and dormancy-like phenomena in other organisms.

## METHODS

### Bacterial strains

Experiments were carried out using *E. coli* EK01 and EK07 strains Krasnopeeva et al. [2019]. The strains have a MG1655 background with the FliC-sticky mutation (EK01), which allows polystyrene beads to stick to bacterial flagella, and the pH sensor pHluorin gene (EK07), both incorporated into the chromosome. For ATP concentration measurements, the EK01 strain was transformed with the plasmid pTP20-QUEEN7* that expresses the QUEEN 7*μ*M* sensor downstream of a strong constitutive promoter as shown in SI Fig. 9. As in Mancini et al. [2020], the plasmid was constructed by PCR amplification of the backbone from plasmid pWR20 Pilizota and Shaevitz [2012] and the sequence containing Q7*. Later steps included purification of the PCR products followed by restriction with the restriction enzymes AvrII and NotI (NEB, UK) and ligation with the T4 DNA ligase (Promega, UK). Chemically competent cells were transformed with the ligation mixes and resulting strains confirmed with colony PCR and sequencing. A map of the plasmid pTP20-QUEEN7* and the primers are given in SI (SI Fig. 8 and SI Table 1). Strains used in this work are listed in (SI Table 2).

### Bacterial growth conditions

For measurements of cellular energetics, cells were grown in M63 (13.6 g/L KH_2_PO_4_, 0.5 mg/L FeSO_4_·7H_2_O, 0.5 g/L MgSO_4_·7H_2_O, 1.27 mg/L Thiamine, 2.64 g/L (NH_4_)2SO_4_ and 0.5% w/v Glucose, final pH: 7.2) to balanced growth at 37°C with shaking at 220 rpm. Balanced growth was achieved by inoculating single colonies into LB medium (0.5% Yeast Extract, 1% Bacto tryptone, 0.5% NaCl) for 3 to 4 h. Cells from such culture were grown to OD 0.3-0.4 in M63 from a starting dilution of 10^−7^. In the case of the EK01-pTP20-Q7* strain, 50 *μ*g/ml of kanamycin was added to the medium.

Optical density readings in the presence of various concentrations of indole, bacteriostatic antibiotics or in carbon or nitrogen deprived medium were obtained in a Spectrostar Omega microplate reader (BMG, Germany) using a flat-bottom 96-well plate that was covered with a lid during the experiments (Costar, UK). Cells grown to balanced growth were collected at OD 0.3-0.4 and transferred to the dormancy-inducing medium at a 1:1 dilution, such that the optical density at the beginning of the measurements was within the microplate readers sensitivity range (~ 0.2). Empty wells adjacent to the samples were filled with water to minimize evaporation. Plates were grown at 37°C with 700 rpm shaking (double orbital mode) for 5 or 24 h with readings taken every 12 min.

For fluorescence intensity comparisons (Fig. 3A and SI Fig. 4A) and for *in vitro* studies on cell lysates we inoculated single colonies in LB medium with 50 *μ*g/ml of kanamycin and grew them overnight at 37°C with shaking at 220 rpm. We then inoculated RDM with glucose Brouwers et al. [2020], University Of Wisconsin (USA) [2020] with 50 *μ*g/ml of kanamycin with the overnight culture at 1:1000 dilution, and grew it at 37°C with shaking at 220 rpm to OD 0.3-0.5.

### Fluorescence microscopy and motor speed measurements

Imaging was carried out in a custom-built microscope with a 100x oil immersion objective lens (Nikon, Japan) Krasnopeeva et al. [2019], Rosko et al. [2017]. Illumination for the cells expressing pHluorin sensor was provided by a neutral white LED with the filter ET470/40x (Chroma Technology, USA) and a UV LED (Cairn, UK). Cells expressing the Q7* sensor were illuminated with the same UV LED and a 488 nm laser (Vortran Laser Technology, USA). Images were taken with an iXon Ultra 897 EMCCD camera (Andor, UK) Krasnopeeva [2018], Rosko et al. [2017]. For both sensors, emission was measured with the ET525/40m filter (Chroma Technology, USA). In the case of pHluorin, images were taken at 1.5 min intervals, exposure time was 50 ms and Andor camera gain 100. Q7* was instead imaged only once per field of view with the same exposure time but with 50 Andor camera gain. Intensity comparisons were also carried out at 50 ms exposure time and 50 Andor camera gain and kept constant across different sensors. Cells were imaged in a custom-built flow-cell (Mancini et al. [2020]), and attached to the coverslip surface as before Mancini et al. [2020]. Briefly, 0.1% Poly-L-Lysine (Sigma, UK) were flushed through the flow cell and washed with 3-5 ml of growth media after 10 s. Cells’ flagellar filaments were sheared as before Krasnopeeva et al. [2019], Mancini et al. [2020], Rosko et al. [2017] by passing them through two syringes with narrow-gauge needles (26 gauge) connected by plastic tubing. 200 *μ*l of cells were delivered to the flow-cell and allowed to attach for 10 min, after which the unattached cells were removed with 1 ml of growth medium. Polystyrene particles (beads) with a diameter of 0.5 *μ*m (Polysciences, USA), were then delivered into the flow-cell and allowed to attach to the filament stubs. After 10 min unattached beads were removed with 1-2 ml of growth media. During measurements, fresh medium was constantly delivered with a peristaltic pump (Fusion 400, Chemyx, USA) using 100 *μ*l/min flow rate. Cells grow attached to the Poly-L-Lysine surface with expected growth rates (given the medium), as previously reported Wang et al. [2019]. Motor speed measurements are carried out at a frequency of 10 kHz (passed through anti-aliasing filters before recording).

### QUEEN sensors characterization from lysates

To characterise the sensors in their original conformation while maintaining conditions as similar as possible to the *in vivo* environment, we performed spectroscopic and microscopic assays on cell lysates. Cells grown in RDM were harvested via centrifugation at 11000g at 4°C for 30 min. Supernatant was discarded and cells resuspended in ice cold QUEEN buffer (50 mM HEPES, 200 mM KCl, 1 mM MgCl_2_, 0.05% Triton X-100, protease inhibitor cocktail (Sigma, GB), pH adjusted to 7.7) Yaginuma et al. [2014]. 100 *μ*ug/ml of lysozyme was added to the cell suspension and incubated for 15 min at room temperature (21°C). Cells were disrupted with a Soniprep 150 sonicator (MSE, UK) with 4 cycles of 30 s on and 30 s off, making sure that no foam was produced. During sonication the tube containing the sample was kept in ice cold water to limit protein damage. The lysates were then filtered with a 0.22 *μ*m filter to remove intact cells and large debris. Solutions with different concentrations of Mg-ATP (Sigma, UK) in QUEEN buffer were mixed with QUEEN lysate maintaining a constant ratio between the two (9:1) in order to maintain a constant protein concentration.

Samples for imaging were prepared in tunnel slides Rosko et al. [2017] by flushing in sequence: 10 *μ*l 0.1% Poly-L-Lysine (Sigma, UK), 100 *μ*l QUEEN buffer wash, 10 *μ*l of a 1 *μ*m polystyrene bead solution (Polysciences, USA), 100 *μ*l QUEEN buffer wash, 10 *μ*l of QUEEN lysate and ATP solution. No incubation was needed between the different steps. The slides were then sealed with CoverGrip*^TM^* sealant (Biotium, US). Three distant (at least 3 fields of view apart) fields of view were imaged with epifluorescence microscopy from each slide, the imaging plane was chosen by focusing on the polystyrene beads.

Fluorescence spectra of QUEEN sensors were obtained with a SPEX Fluoromax 3 spectrometer (Horiba, JP). Excitation was scanned from 370 to 500 nm at 513 nm emission with a slit size of 3 nm. The temperature of the sample holder was controlled via a circulating water bath and readings were taken after letting the temperature of the sample reach equilibrium with that of the sample holder (24°C), i.e. after at least 5 min.

### MIC estimation

MIC was estimated on EK07 cells grown to balanced growth. Aliquots of cultures in balanced growth were diluted 1:100 into pre-warmed M63 medium (final OD 0.002-0.004) with different concentrations of ciprofloxacin (2, 4, 8, 16, 32, 64, 128, 256 ng/ml) or kanamycin (2, 4, 8, 16, 32, 64, 128, 256 *μ*g/ml). Cell suspensions were then transferred into a pre-warmed 96-well plate, incubated for 24 h and imaged (SI Fig. 8).

### CFU counting

For CFU counting cells were grown to balanced growth, harvested via centrifugation at 8000g for 5 min and washed twice in the various dormancy-inducing conditions or fresh M63. Cultures were then incubated at 37°C with 220 rpm shaking (the same conditions in which they had been grown). From this point onward the protocol varies slightly depending on whether we are trying to estimate survival to dormancy-inducing conditions (Fig. 1B) or to the combination of dormancy-inducing conditions and antibiotic treatments (Fig. 4).

For estimating the number of survivors to our dormancy-inducing conditions (Fig. 1B), aliquots of the cultures were taken after 60, 120, 180 and 240 min (Fig. 4A), pelleted at 8000g for 2 min and washed twice in fresh M63 medium. Samples were then serially diluted with a 1:9 ratio in fresh M63 and 10 *μ*l of each dilution spotted on rectangular petri dishes (Thermo Fisher Scientific, UK) containing LB agar. The petri dishes were next tilted to let the liquid spread across the plate as in Jett et al. [1997], incubated for 24 h at 37° C and colonies counted.

For estimating survival to antibiotics (Fig. 4), cells were left in the dormancy-inducing conditions for 30 min. Once dormancy had been established, antibiotics at 5x the MIC (16 ng/ml for ciprofloxacin and 8 *μ*g/ml for kanamycin (SI Fig. 8)) were added to the cultures. Aliquots of the cultures were taken after 30, 120, 180 and 240 min and plated as explained above (Fig. 4A).

Experiments were carried out in biological triplicates.

### Data analysis

#### Motor speed traces

Raw traces of the x and y position of the bead attached to the filament stub were analyzed by a moving-window discrete Fourier transform in LabView as in Rosko et al. [2017]. From the obtained motor speed traces DC frequency (50 Hz) was removed, speeds lower then 5 Hz ignored, and subsequently a median filter (window size 31) was applied Krasnopeeva et al. [2019], Mancini et al. [2020]. We use a chemotactic wild type strain for which the flagellar motor can change rotational direction, which appears as a negative speed after application of the moving-window Fourier transform. For the purpose of the PMF measurements these short intervals can be disregarded, and we only show the speed values above 0 Hz as in Mancini et al. [2020]. Because of our flow slide design Mancini et al. [2020], media switches during treatments happen 2 to 4 min from the start of the recordings. To account for this variability, traces of cells exposed to carbon starvation were aligned taking as a reference an arbitrary motor speed value of 150 Hz, which in our conditions is a good indicator of the timing of bacterial flagellar motor speed drop. For all traces in Fig. 2C, PMF values are normalized to the initial PMF calculated as the average of the first 60 seconds of the trace.

#### Fluorescence images

The image analysis was carried out with a custom written PYTHON script. Fluorescence images were initially inspected for any unevenness in the epifluorescence illumination (i.e in the the field of view) by assaying background greyvalues using the straight line tool of ImageJ Schneider et al. [2012]. To individuate the cells, objects with high grey values were discerned from the background by applying a global threshold via the Otsu’s method Otsu [1979] and labeled. All found objects were inspected for size and aspect ratio so that anything smaller than 8 *μ*m^2^ or bigger than 56 *μ*m^2^ or with an aspect ratio below 1.7 were excluded. The criteria were set based on the average size, length and width of cells in our growth condition (balanced exponential growth at the growth rate of 0.7 h^−1^) and allow us to discard labels that are not correctly segmented and small artifacts. Total cells’ intensity values were obtained by summing up and averaging pixel intensities of selected objects.

#### ATP calibration curves

The sensitivity range of the Q7* protein is independent from imaging conditions and settings. Differences in the optical setup that can alter light intensity, even those arising from small changes in the alignment of the optics, will however change the absolute value of the ATP to 395/488nm relationship. Calibrations therefore need to be performed periodically (or after any known changes in optics). For comparison, in Fig. 3B we eliminated the variability between calibrations performed in different days by rescaling the values to the range of the curve used to calculate ATP concentrations *in vivo* in Fig. 3D. We used the following formula: 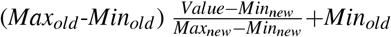.

#### Plate reader data

Curves from different repeats were averaged and the standard deviation estimated. Experiments carried out on different days can have slightly different lag times due to small differences in preparation times. To make the results better visible we omitted lag periods where these were slightly longer but did not align the data in any other way.

#### Spectroscopy

The intensity of the light source of the SPEX Fluoromax 3 spectrometer (Horiba, JP) across the range of excitation wavelength examined was measured during data acquisition and each fluorescence measurement obtained normalised by it, thus eliminating differences due to the light source. The results of the experiments, which were carried out in triplicate, were averaged and the standard deviation calculated. The spectra obtained were normalised by the average of the replicates’ values at 435 nm excitation, which is the isosbestic point of the QUEEN spectrum, insensitive to ATP concentration.

## AUTHOR CONTRIBUTIONS

L.M. and T.P. conceived the experiments. L.M. performed the experiments and analyzed experimental data. L.M. and T.P. interpreted the results and wrote the manuscript.

## ACKNOWLEDGEMENTS

We are grateful to Hideyuki Yaginuma for donating us constructs containing the QUEEN sensors and for providing us the published data with QUEEN excitation spectrum. We are also grateful to Alessia Lepore and Meriem El Karoui for donating us an aliquot of ciprofloxacin and to members of Pilizota lab as well as Alessia Lepore, Meriem El Karoui, Ssu-Yuan Lin, Chien-Jung Lo, Thomas Julou for useful discussions. We are grateful to Pietro Cicuta for advice and support during the preparation of the manuscript.

This work was financially supported by the Cunningham Trust scholarship ACC/KWF/PhD1 to T.P. and L.M.. T.P. acknowledges the support of Human Frontier Science Program Grant (RGP0041/2015), and UK Research Councils Synthetic Biology for Growth program and is a member of the Biotechnology and Biological Sciences Research Council/Engineering and Physical Sciences Research Council/Medical Research Council-funded Synthetic Biology Research Centre (BB/M018040/1).

## DATA AND MATERIALS AVAILABILITY STATEMENT

All data is deposited at (link will be provided upon acceptance). Materials are available upon request to the corresponding author.

