## Supplementary information for "Environmental conditions define the energetics of bacterial dormancy and its antibiotic susceptibility"

### SUPPLEMENTARY FIGURES

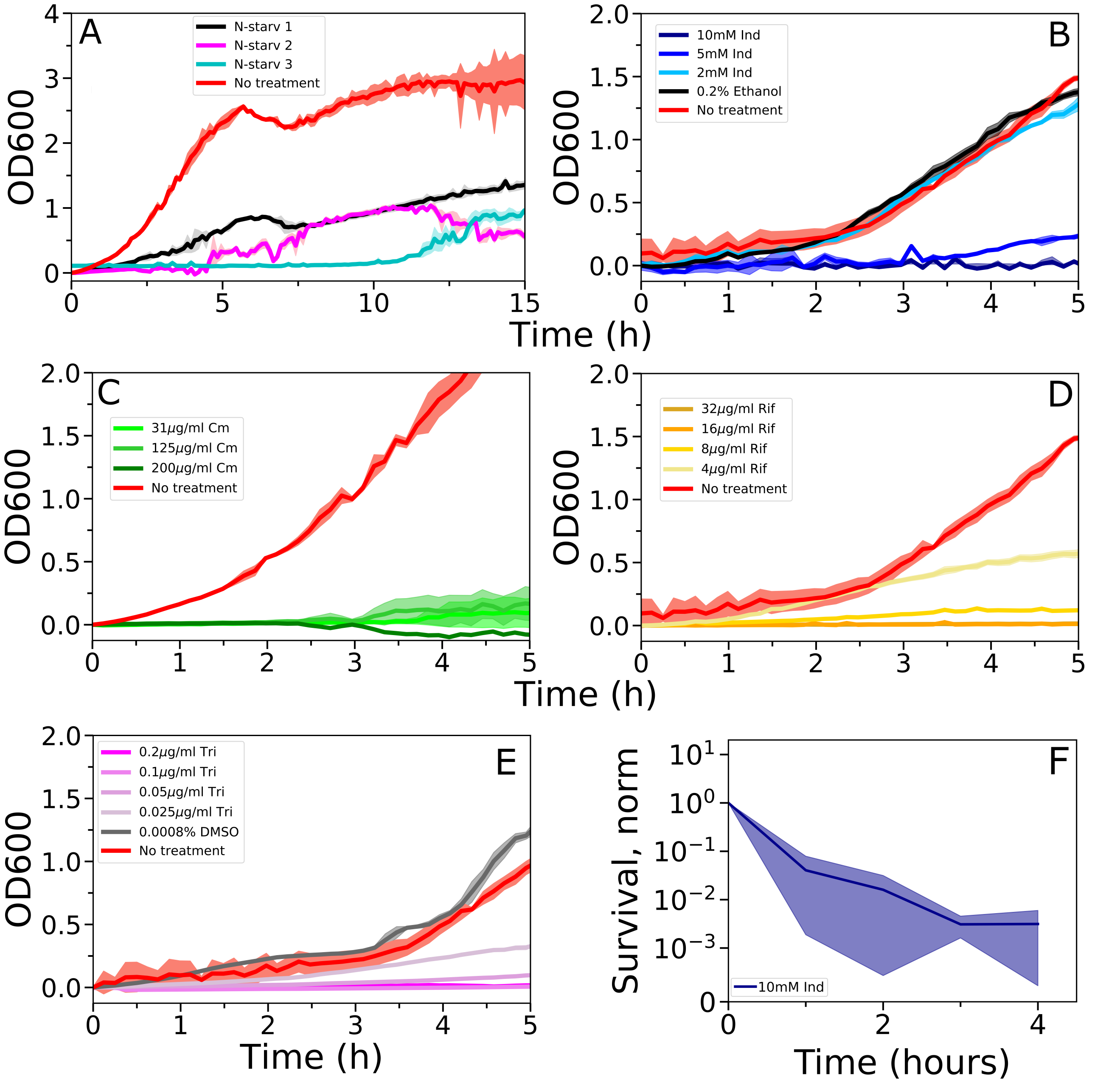


Supplementary figure 1: Individuation of dormancy inducing conditions by monitoring cell growth in a plate reader. OD of an *E. coli* culture immediately after **a,** transfer into M63 deprived of the nitrogen source (ammonium sulphate), addition of **b,** 10, 5, 2 mM indole or 0.2% ethanol (ethanol is the solvent of indole and 0.2 % is its maximum concentration at 10mM indole), **c,** 0, 31, 125 and 200 $\mu$M chloramphenicol, **d,** 1, 2, 4, 8, 16, 32, 64 and 128 /*mu*M rifampicin, **e,** 0.025, 0.05, 0.1 and 0.2 $\mu$g/ml trimethoprim or 0.0008% DMSO (solvent of trimethoprim and 0.0008% is its maximum concentration at 0.2 $\mu$g/ml trimethoprim. Traces indicate mean and standard error (shaded area) from three replicates. Negative control in **a** and **c** is re-plotted from Fig. 1A of the main text. In **a,** results from three independent experiments carried out in triplicates are given to illustrate the variability likely linked to different amounts of nitrogen traces present in the medium. **f,**10 mM indole stops growth but does not preserve cell viability. Survivors are estimated using the CFU counting method as explained in the main text.

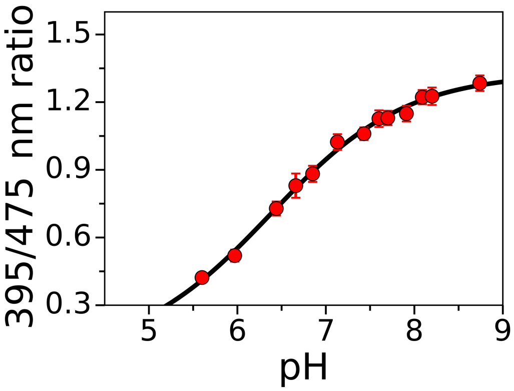


Supplementary figure 2: Calibration of the pHluorin sensor performed *in vivo* at 37°C in *E. coli* cells treated with a mixture of 40 mM potassium benzoate and 40 mM methylamine hydrochloride (PBMH) which permeates the cell membrane and neutralizes ΔpH. External pH was varied by controlling the pH of the buffer with KOH or HCl Krasnopeeva et al. [2019], Wang et al. [2019]. Each point is the average of $\sim$ 50-250 cells. Error bars show the standard deviation.

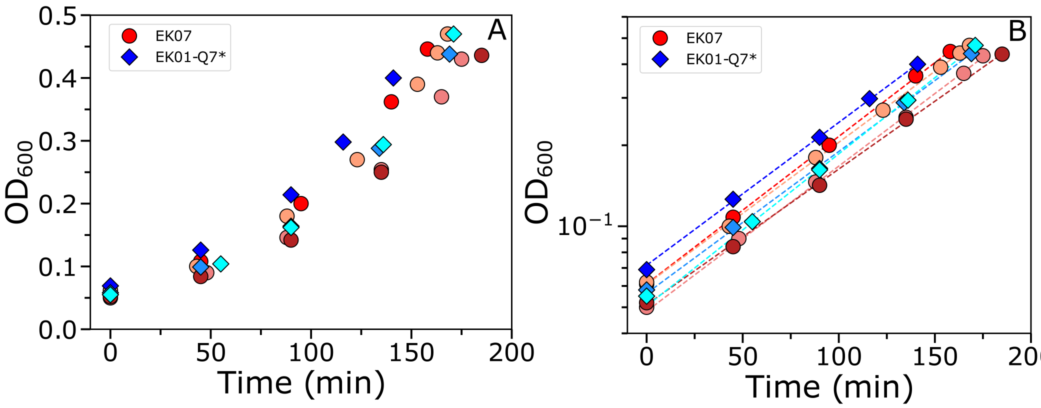


Supplementary figure 3: The two *E. coli* strains we use, EK07 and EK01-Q7*, have similar growth rates when cultured in M63 + Glucose. **a,** Growth curves for the strains EK07 (circles) and EK01-Q7* (diamonds) measured by optical density readings performed every 45 min in a spectrophotomer. Different shades of red and blue indicate independent replicates for EK07 (red, 4 replicates) and EK01-Q7* (blue, 3 replicates). **b,** Data points from **a** presented on a log scale. Dashed lines are exponential fits to the data for each replicate. The slope of the fits is the exponential growth rate and corresponds to (0.73±0.03) h$^{-1}$ for EK07 and (0.74±0.02) h$^{-1}$for EK01-Q7* (calculated as average of the three measurements for each condition), which is in line with previous measurements for the medium Scott et al. [2010].

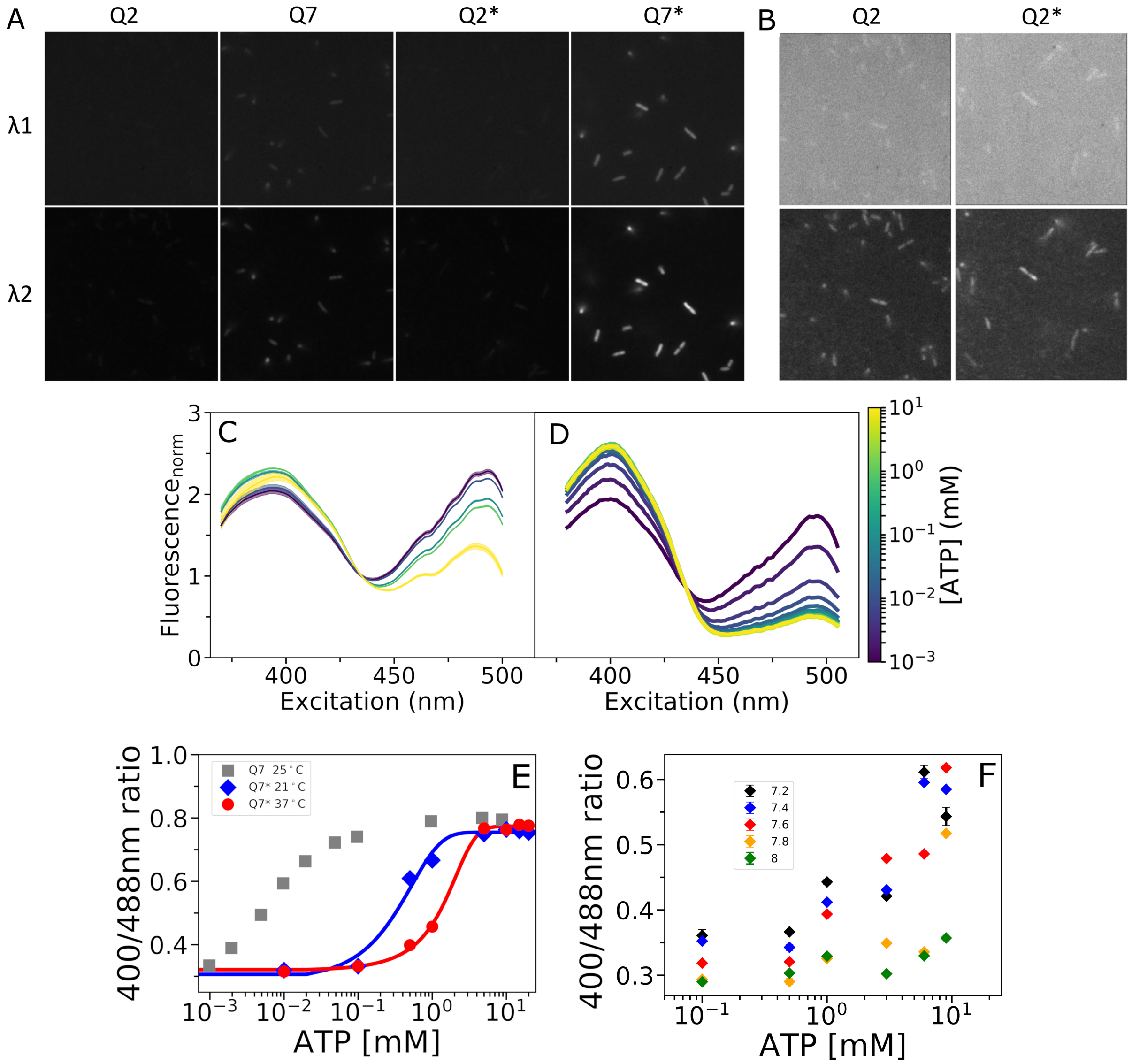


Supplementary figure 4: Characterization of the Q7* sensor. **a,** Example fields of view of *E. coli* MG1655 expressing QUEEN 2mM (Q2) and QUEEN 7$\mu$M (Q7) from Yaginuma et al. [2014], and the same sensors without the histidine tag (Q2*) and (Q7*) obtained with fluorescence microscopy at two different wavelengths. The illumination settings for the different strains were kept fixed at 50 ms exposure time, and 50 EMCCD camera gain. **b,** Q2 and Q2* imaged with the same settings of **a,** but the greyvalue ranges of the images are rescaled (and kept constant between the two) to make them visible and comparable. Cells were grown to balanced exponential growth in RDM with glucose (see the *Methods* in the main text) and imaged in a tunnel slide (design given in Rosko et al. [2017]).

Supplementary figure 4: **c,** The fluorescence excitation spectrum of lysates obtained from cells expressing the QUEEN 7$\mu$M* sensor (see main text *Methods*) assayed at 513 nm emission. Colours indicate different ATP concentrations as shown in the colour bar on the right (10, 1, 0.1, 0.01 and 0.001 mM). Autofluorescence signal of the wild type is subtracted from each curve. For each curve values were normalised to the value at 435 nm, which is insensitive to ATP concentration (Yaginuma et al. [2014]). Each trace shows the mean of three measurements and the standard deviation (shaded area). The experiment was carried out at 24°C. **d,** Spectra of Q7 at 25°C re-plotted from Yaginuma et al. [2014] as a comparison. As in **c**, we normalized spectra by the fluorescence value at 435 nm. The differences in ratios are a consequence of different experimental conditions and preparations of the fluorescent samples. In Yaginuma et al. [2014], spectroscopic analysis was carried out on the purified protein, as opposed to our cell lysates. **e,** The sensitivity range of the QUEEN 7$\mu$M* sensor shows a temperature dependence (blue diamonds, red circles). As a comparison, the Q7 sensor calibration at 25°C from Yaginuma et al. [2014] is re-plotted in grey, as a comparison, where the data was re-scaled using $y_{res} = (0.8-0.3)(y-y_{min})/(y_{max}-y_{min})+0.3$, where $y$, $y_{res}$, $y_{min}$ and $y_{max}$ are the original, the rescaled, the minimum and the maximum value of the ratio, respectively. The procedure was performed to enable comparison of the ATP sensitivity range. For the Q7* data, error bars representing the standard deviation for three measurements are smaller than the markers, which indicate the mean. As before, the calibration for Q7* was obtained from cell lysates. **f,** The calibration curves taken at different pH values show QUEEN 7$\mu$M* sensor pH sensitivity. Symbols show the mean and error bars, where visible, the standard deviation.

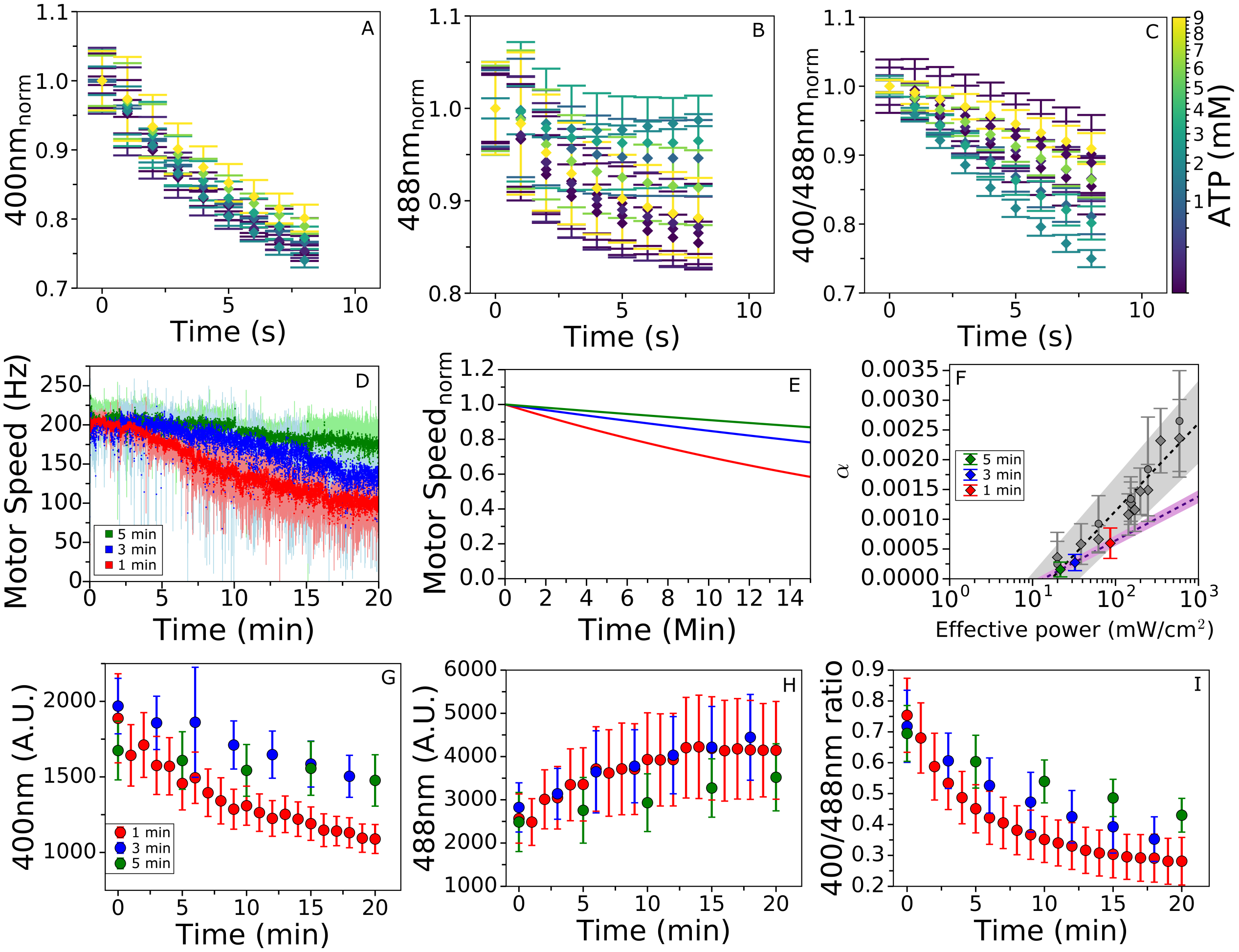


Supplementary figure 5: Defining *in vivo* imaging conditions for ATP measurements with the QUEEN 7$\mu$M* sensor. **a-c** *In vitro* photoeffects of QUEEN 7$\mu$M*. Normalized intensity of QUEEN 7$\mu$M* lysates observed under 400 nm illumination in **a** and 488 nm illumination in **b**, and at different ATP concentrations (as indicated by the colour map to the right). **c,** gives the ratio of the two. Data points are averages of fluorescent values obtained from 3 different fields of view. The error bars show the standard deviation. Values were normalized by the signal intensity at time t = 0. **d,** PMF loss due to photodamage is monitored via the flagellar motor speed while exposing the cells to illumination at 400 and 488 nm for 20 min at different frame rates. Averages of 3 cells from 3 different experiments per condition are given. Shaded areas show the standard deviation. **e,** Light damage causes a power-dependent (effective power that takes into account the LED power, exposure time, duration of recording and frequency of recording) exponential decay in the flagellar motor speed Krasnopeeva et al. [2019]. To quantify the damage, traces in **d** are normalized by their value at time t = 0 and fitted to a single exponential, $y = e^{-\alpha x}$ Krasnopeeva et al. [2019]. From the fit we obtained $\alpha$ equal to 0.0006, 0.00027 and 0.00016 for 1,3 and 5 min intervals between illumination events, respectively. **f,** $\alpha$ parameters from the fits in **e** plotted against the effective power, calculated as in Krasnopeeva et al. [2019]. In grey we re-plot the results from Krasnopeeva et al. [2019]. The three $\alpha$ values obtained in **e** are fitted to a logarithmic function $a \cdot logP_{eff} + b$, as in Krasnopeeva et al. [2019]. $a$=0.00064 and $b$=-0.00181 for the previously published data and $a$=0.00032064 and $b$= -0.000835 for the data obtained here. The shaded area shows the standard deviation of the fit. **g, h** and **i** Fluorescence signal from cells treated with 5% ethanol, when illumination is administered at 1 (red), 3 (blue) and 5 (green) min intervals. Error bars show the standard deviation and each time point was obtained from 15 cells or more. **g** shows the signal obtained from 400 nm excitation illumination, **h** from a 488 nm excitation and **i** the ratio of the two.

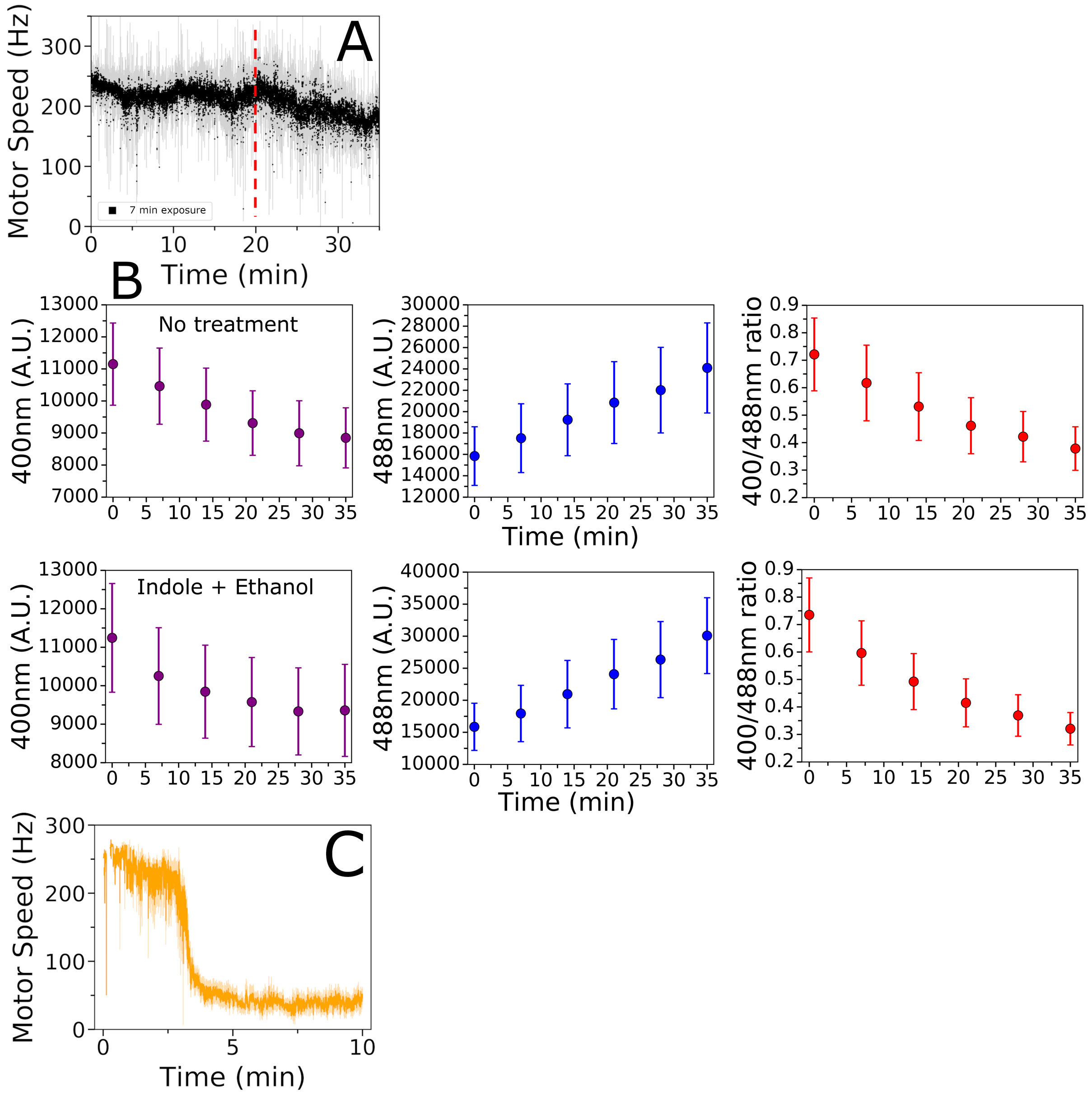


Supplementary figure 6: Photoeffects limit the use of QUEEN for time series measurements. **a,** PMF of untreated *E. coli* measured via the flagellar motor speed over 35 min, showing that the PMF does not change over a 20 min period (red dashed line) when cells are illuminated every 7 min with 400 and 488 nm wavelength with 50 ms exposure times. Average trace of 5 single cells is shown with shaded area giving the standard deviation. **b,** Average 400/488 nm fluorescence intensity ratio (red), and fluorescence intensity obtained at 400 nm (purple) and 488 nm (blue) excitation from QUEEN 7$\mu$M* sensor in: untreated cells (M63) and cells exposed to 2% ethanol and 10 mM indole (in M63). The error bars show the standard deviation from $\sim$50 cells. **c,** Motor speed trace of cells treated with a mixture of 10 mM indole and 2% ethanol which rapidly shut down PMF. (The cells are exposed to the mixture after 2-3 min from the beginning of the recording). The trace is the average of 10 cells from 10 independent experiments. The shaded area indicates the standard error.

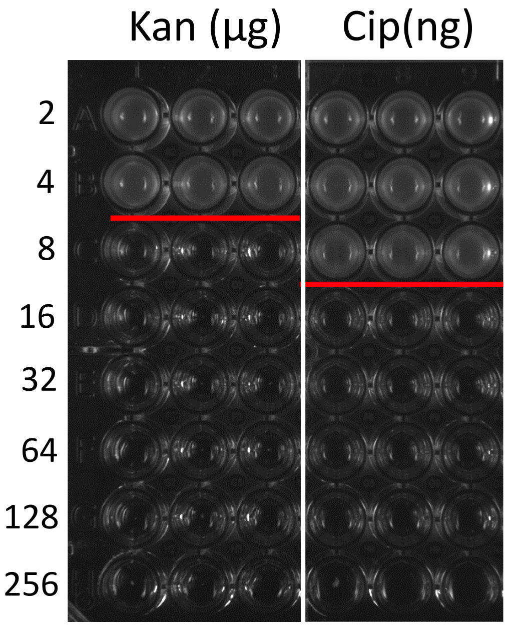


Supplementary figure 7: We estimated MIC for EK07 strain treated with kanamycin or ciprofloxacin antibiotics. The assay was carried out in triplicates (columns show experimental repeats). The rows indicate antibiotic concentration in $\mu$g/ml for kanamycin and in ng/ml for ciprofloxacin. The estimated MICs are shown by the red line: 8 $\mu$g/ml for kanamycin and 16 ng/ml for ciprofloxacin. Cells grown to balanced growth were inoculated in media supplemented with the various concentrations of antibiotic at a 1:100 dilution, with a final OD of 0.002-0.004.

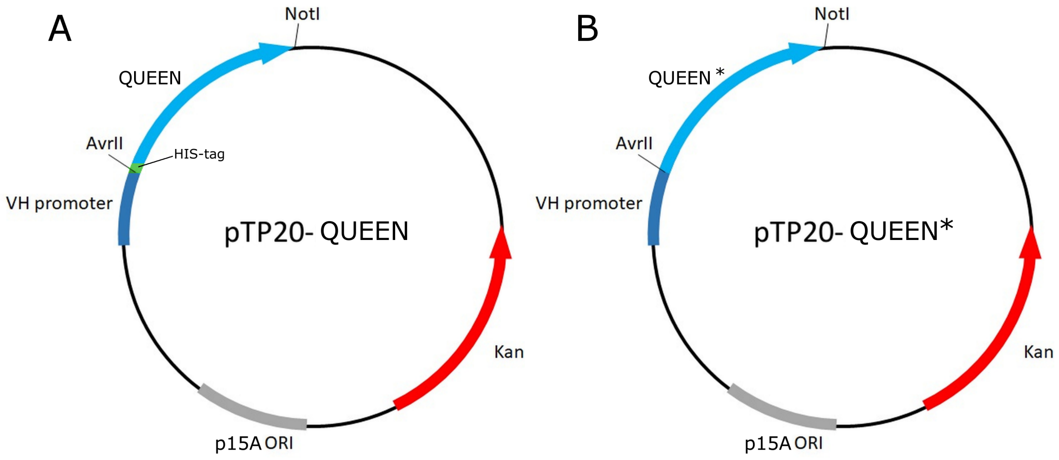


Supplementary figure 8: Plasmid maps of **a,** pTP20-QUEEN (Q2 and Q7) and **b,** pTP20-QUEEN* (Q2* and Q7*) constructed from pWR20 plasmid [Pilizota and Shaevitz, 2012]. The insertion site of the sensor, constitutive cytochrome oxidase promoter from *Vibrio Harveyi*, the origin of replication, the Kanamycin resistance cassette and the position of the His-tag for QUEEN are shown.

### SUPPLEMENTARY TABLES

| Plasmid | Fragment | Template | Primers |
|---|---|---|---|
| pTP20-Q2* | pWR20 backbone | pWR20 | 5' AAAGCGGCCGCGGTGATTGATTGAGCAAG 3'<br>5' AAACCTAGGATGTATATCTCCTTAACTAGGT 3' |
| | Q2* | Q2mM | 5' ATACCTAGGATGAAAACTGTGAAAGTGAAT 3'<br>5' AATGCGGCCGCTCACTTCATTTCCGCAAC 3' |
| pTP20-Q7* | pWR20 backbone | pWR20 | 5' AAAGCGGCCGCGGTGATTGATTGAGCAAG 3'<br>5' AAACCTAGGATGTATATCTCCTTAACTAGGT 3' |
| | Q7* | Q7$\mu$M | 5' ATACCTAGGATGAAAACGATCCACGTGA 3'<br>5' AATGCGGCCGCTCACTTCATTTCCGCAAC 3' |
| pTP20-Q2 | pWR20 backbone | pWR20 | 5' AAAGCGGCCGCGGTGATTGATTGAGCAAG 3'<br>5' AAACCTAGGATGTATATCTCCTTAACTAGGT 3' |
| | Q2 | Q2mM | 5' ATACCTAGGATGCATCACCACCATCATCACAAAA CTGTGAAAGTGAATAT 3'<br>5' AATGCGGCCGCTCACTTCATTTCCGCAAC 3' |
| pTP20-Q7 | pWR20 backbone | pWR20 | 5' AAAGCGGCCGCGGTGATTGATTGAGCAAG 3'<br>5' AAACCTAGGATGTATATCTCCTTAACTAGGT 3' |
| | Q7 | Q7$\mu$M | 5' ATACCTAGGATGCATCACCACCATCATCAC AAAACGATCCACGTGAGC 3'<br>5' AATGCGGCCGCTCACTTCATTTCCGCAAC 3' |

Table 1: Plasmids constructed by restriction-ligation for the experiments shown in this chapter. Fragments and primers are indicated. The pWR20 backbone was obtained from Pilizota and Shaevitz [2012] and carries the p15A Selzer et al. [1983] origin of replication and a kanamycin marker. The QUEEN sensors templates were obtained from Yaginuma et al. [2014].

| Strain | Origin | Figure |
|---|---|---|
| EK01 | Krasnopeeva et al. [2019] | 3A,SI5A,SI5C,SI5D |
| EK07 | Krasnopeeva et al. [2019] | 1,2,3B,4,SI1,SI2,SI3,SI4,SI9 |
| EK01-pTP20-Q2 | This work | 3A,SI5A |
| EK01-pTP20-Q7 | This work | 3A,SI5A |
| EK01-pTP20-Q2* | This work | 3A,SI5A |
| EK01-pTP20-Q7* | This work | 3A,3B,SI3,SI5A,SI5B,SI5D,SI5E,SI6,SI7,SI8 |

Table 2: List of *E. coli* strains used in this work.
